# A mathematical model for inflammation and demyelination in multiple sclerosis

**DOI:** 10.1101/2025.07.31.667831

**Authors:** Adrianne L. Jenner, Georgia R. Weatherley, Federico Frascoli

## Abstract

Multiple sclerosis (MS) is an incurable, life-long disease caused by the demyelination of neurons in the brain and spine. MS is often characterised by relapses in inflammation and demyelination, that are then followed by periods of remittance. Symptoms can be highly debilitating and there are still many open questions about the origin and progression of the disease. Mathematical modelling is well-placed to capture the dynamics of MS and provide insight into disease aetiology. In this work, we present a minimal model for MS disease onset and progression driven by inflammation and demyelination. The model dynamics are capable of describing a typical evolution of the illness, with changes from a healthy state to a diseased scenario captured by certain ranges of parameter values. Our model also describes the non-uniform oscillatory nature of the disease, born from a Hopf bifurcation due to the strength of the inflammatory response. In particular, using experimental data for Contrast Enhancing Lesions (CELs) obtained from MS patients, we are able to reproduce some of the typical relapsing-remitting behaviours of this disease. We hope that the model presented here can serve as a baseline for more complex approaches and as a tool to predict possible evolutions of the disease.

## 1. INTRODUCTION

Multiple sclerosis (MS) is a neurodegenerative disease caused by inflammatory events that result in the demyelination of axons, leading to a range of physical and cognitive symptoms. With the increasing incidence of MS, particularly in women (20), there is an urgent need to improve our understanding of the aetiology and treatment of this condition, especially in a quantitative and systematic way.

While the immune involvement in MS is complex, the disease progression is largely driven by antigen-presenting cells (APCs). These cells present myelin antigen to lymphocytes, which then in turn cause damage to healthy myelin surrounding neurons (1). Apart from APCs, the main immune cells playing a role in MS are B cells and T cells. T cells are often thought to play a helper, regulatory or effector function and can cause demyelination of neurons (38). B cells have been shown to cause damage to oligodendrocytes, the maker of myelin (1). Recent, compelling findings, have implicated Epstein-Barr virus as the trigger for the development of MS, however, questions still remain on how a virus with tropism for B cells causes the development of a disease of the central nervous system (37).

Relapses, sudden worsening or emergence of symptoms (23), are related to focal infiltration of lymphocytes into the brain and spinal cord causing damage to myelin and axons (8; 25). Different types of MS exist and a patient’s history of relapsing activity is used in conjunction with their accumulation of disability to distinguish four disease phenotypes (31), see Fig. 1(A). Relapsing-remitting MS (RRMS) patients experience intermittent relapses amongst prolonged periods of remission. These patients eventually enter secondary-progressive MS (SPMS), undergoing continual neurological decline even though active inflammatory events that characterise the early RRMS stage of disease are no longer present. Primary-progressive MS (PPMS) patients experience a different course and show gradual, persistent worsening from the onset of the disease, with progressive–relapsing MS (PRMS) patients also affected by relapse events (47). Eighty percent of patients initially follow a RRMS disease course (43), experiencing a mean annual relapse rate of around one half (e.g. 0.54) (47). Given its incidence and occurrence, RRMS is the focus of the modelling proposed in this work.

**Figure 1.**
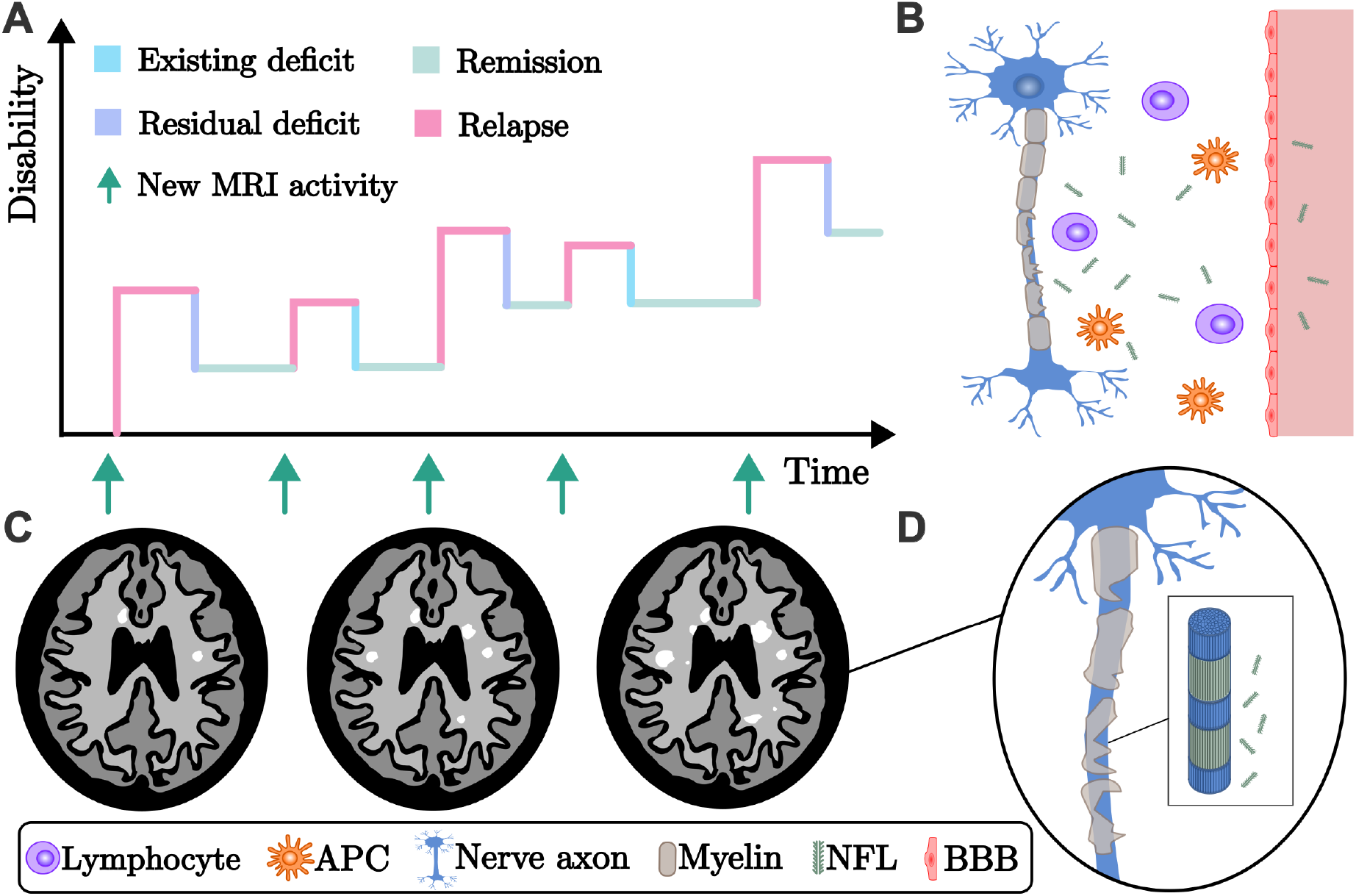
Crucial components of MS disease informing the mathematical modelling. (A) Typical evolution of the EDSS score with time, showing periods of increased disability interspersed with periods of no changes. (B): Pictorial of the phisiology of demyelination, with different immune cells (lymphocytes and antigen presenting cells (APCs)) participating in the process of demyelination and the production of neurofilament light chains (NFLs) that diffuse into the bloodstream, crossing the blood-brain barrier (BBB). (C) Schematic examples of contrast enhancing lesions (CELs) appearing in a typical patient’s MRIs, with demyelinated regions in white. (D) Magnification of demyelination on a neuron, resulting in a damaged axon and NFLs in the bloodstream.

Understanding the factors that contribute to the frequency of relapses in patients with RRMS is crucial, as nearly half of these episodes lead to lasting disability (30). While the exact mechanisms governing relapse frequency remain unknown (23), there are a range of potential relapse triggers, each holding implications for frequency. For example, infections are associated with an increased risk of relapse (25; 47), whereas pregnancy decreases the risk of relapse, particularly during the third trimester (9; 21; 23). Further suspected risk factors include smoking, age, stress levels, ethnicity and vitamin D levels.

Mathematical modelling is well-positioned to support hypothesis generation and mechanistic understanding of MS, although existing models are limited (48). In the models that do exist, the degrees of freedom are often large and, due to the difficulty of performing *in vivo* measurements in the central nervous system (CNS), parameterisation is cumbersome or limited. Despite this, there has been a recent increase in MS mathematical and computational modelling (see, for instance (36; 19; 3; 34; 40; 41; 4)). There are a number of ordinary differential equation (ODE) models of MS that consider changes in immune responses over time (5; 50; 51; 46; 32; 16; 24; 33). These models are concerned with the interaction of damaging immune populations with diseased and healthy elements of the brain, with some including cross-regulatory immune dynamics to understand possible drivers of the oscillatory nature of the disease (32; 46; 24).

Stability analysis in MS modelling has helped to link the oscillatory dynamics that arise from ODE models or PDE models to characteristics of MS disease (10; 12). For instance, Zhang et al. (50) and Zhang and Yu (51) model the regulatory T cell interactions with APCs occurring through a system of ODEs in MS. They emphasise the importance of a Hopf bifurcation as a condition for recurrent behaviour in their models. Elettreby and Ahmed (16) propose a simple model capturing MS dynamics through healthy brain cells, infected brain cells and harmful effector entities (immune cells of virions). They conduct a stability analysis to identify the bifurcation conditions required for recurrence and oscillations in the ODE system they propose. Whilst other non-trivial mathematical approaches focused on oscillatory dynamics do exist (24; 49), they are certainly not minimal and, as such, have not been analysed thoroughly. A recent spatial model by Bisi *et al*.(4) observed spatial-temporal oscillations and pattern formation of lesions captured elegantly through immune interactions between an antigen-presenting population, microglia and immunosuppressive cells. Models, such as these, highlight simple ways in which the complex immunology of MS can be captured at the cellular level without undue complexity, see Fig. 1(B). The challenge remains, however, since measurements at this level are not feasible, and so mathematical models that can be linked to available clinical data at the symptom or MRI level are needed.

MS is often characterised by highly irregular and noisy dynamics of peaks and relapses that can be seen in the Expanded Disability Status Scale (EDSS) and contrast-enhancing lesions (CELs) obtained from Magnetic Resonance Imaging (MRI), see Fig. 1(C). MRI techniques detect local inflammatory events on T1-weighted images to visualise and quantify damage (6). In contrast, the EDSS measures the level of disability experienced by a patient over time by assessing changes in functionality (26). These two measurements are the main clinical scores used to quantify disease stage and progression.

In this work, we present the simplest possible two-state model possible of MS inflammation and demyelination, conducting an in-depth analysis to reveal the existence and characteristics of oscillations predicted in neuronal myelin density. We show, for example, how the model adequately describes CEL measurements and how the interplay between myelination levels and inflammation can explain the development of MS in typical circumstances. The equations can also provide idealised scenarios for patients’ prognosis. The paper is organised as follows: in Section 2 we introduce the model and describe its biological and medical meaning, in Section 3 a local stability analysis is performed to show the model’s typical equilibria and dynamics, in Section 4 a bifurcation analysis shows the range of dynamics that the model can sustain and, finally, in Section 5, a comparison with real patient data is proposed, showing the model’s capability to reproduce actual data from CELs. The paper ends with the Conclusions Section, where a general discussion of the findings and limitations of our approach is conducted.

## 2. DESCRIPTION OF THE MATHEMATICAL MODEL

For the minimal model proposed here, we consider two populations: *M* (*t*) represents the proportion of viable healthy myelin and *I*(*t*) describes the level of inflammation. Spatial effects are ignored and we assume a “well-mixed” system that captures what occurs “on average” spatially over time. We adopt a pedagogical point of view where the variables and parameters of the system are representatives, or approximations, of more complex, often not completely understood, physiological and pathological phenomena.

The full system of ODEs modelling MS disease corresponding to Fig. 2(A) is given below by:

**Figure 2.**
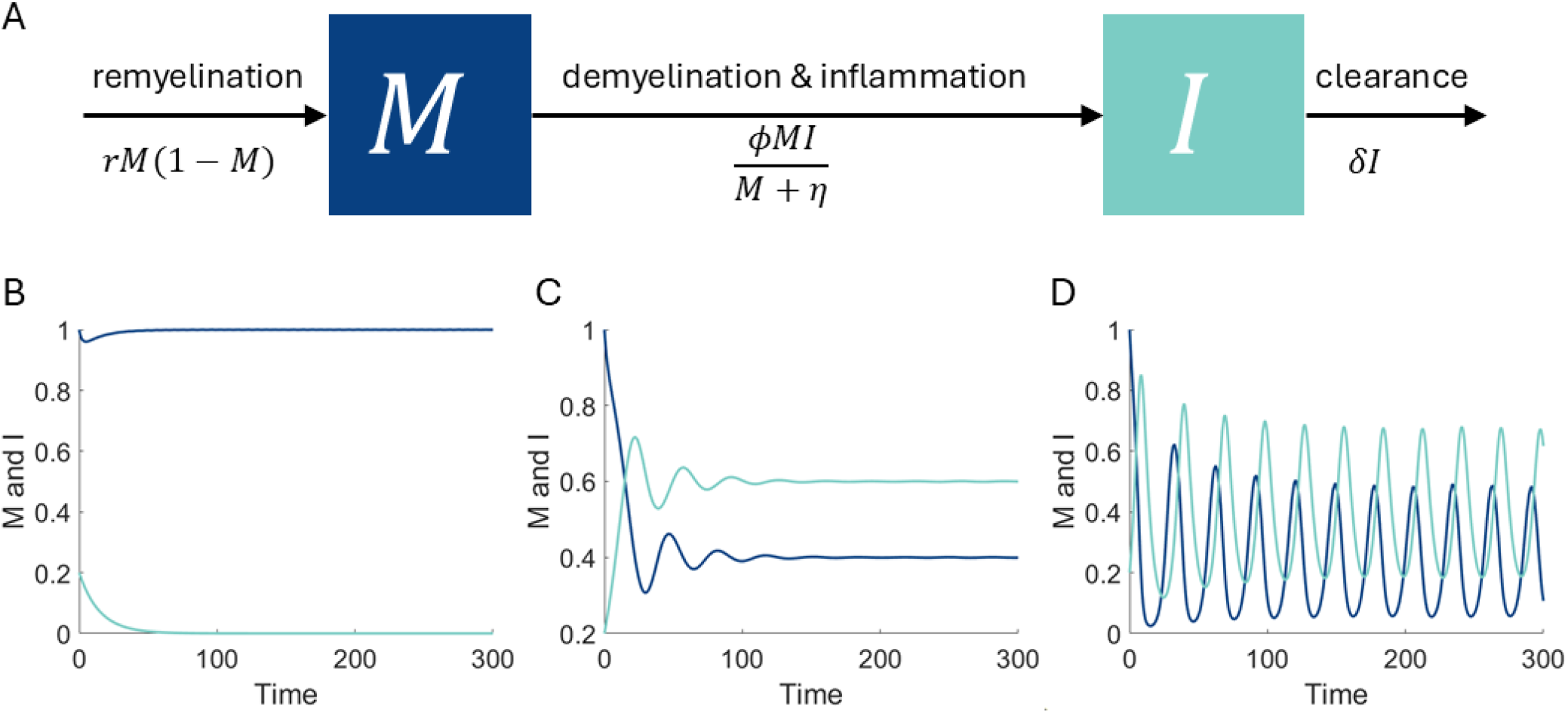
Dynamics of the model for MS in Eq. 1. (A) Compartment schematic describing remyelination and loss of myelin *M* (*t*), as well as the increase and decrease of inflammation *I*(*t*). (B)-(D) Simulations of the model for *r* = 0.5,*δ* = 0.2, and *η* = 0.5 where (B) *ϕ* = 0.2, (C) *ϕ* = 0.45 and (D) *ϕ* = 0.7. (B) The model shows robustness to inflammation, which causes only a small, brief decrease in myelin and the system eventually returns to the healthy state (*M* ^(1)^, *I*^(0)^). (C) The system shows transient oscillations that gradually converge to the diseased state (*M* ^(*\**)^, *I*^(*\**)^). (D) The model predicts stable period oscillations with a period of *t* = 30, capturing relapse and remittance in MS patients.

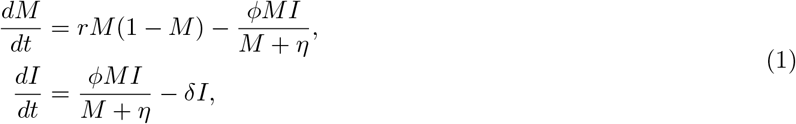

where *t* is time. Decreases in *M* from unity indicate proportion of myelin lost in areas of the CNS. This phenomenon is highly heterogeneous, with lesion densities that vary according to the part of the CNS and the individual characteristics of the patient (42). Axonal loss, for example, can attain densities of 20% – 30% on average, compared to white matter control subjects (45). As such, *M* is assumed to describe an “average” loss that could occur in different parts of the patient’s CNS. We assume all relevant biological dynamics occur for *r, ϕ, η* and *δ >* 0 as setting any of these parameters to zero removes a major assumption in the model.

In contrast, an increase in *I* represents an increase in inflammation. Identifying a single biomarker for *I* is inherently complex; however, neurofilament light chains (NFLs) represent the most reliable proxy (44), see Fig. 1(D). Neurofilaments are the single most abundant protein in myelinated axons. Physiological turnover and damage to neurons is thought to be the origin of neurofilaments measurable in the cerebrospinal fluid and blood (44). NFLs is a subunit of the neurofilament protein and a higher presence of NFLs in blood serum is correlated with neuroaxonal injury (18). Increases in the level of NFLs, between 0.2 and 2-3 times baseline, have been found in the serum of MS patients (14; 13; 39). We, therefore, assume that *I* reflects the increase of NFLs’ presence in the bloodstream and that its value is correlated to NFL’s density in the serum of MS patients.

Rates *M* and *I* vary with time *t*, where we assume that a unit time (*t* = 1) equals one month. Parameter *r* captures the rate of remyelination, which is usually on the order of several months. It is worth noting that, in many MS cases, remyelination is never thoroughly complete (7; 29). Since myelin is modelled as regrowing logistically (see first term of Eq. 1), a value of *r* between 0.3 *−* 1.0 is chosen: this implies that a ratio of repair of about 90% (i.e. *M* = 0.9) takes between two and seven months to occur, in the absence of further inflammation and with an initial myelination rate *r* = 0.5. The parameter *δ* is the rate of decay of inflammation. Usually, the average duration of an inflammatory episode is on the order of about a month or so, with some examples of inflammation that last for several months (42), so *δ* is chosen around unity. The value of *η* represents the critical value of myelin at which the demyelination process and activation of the inflammatory immune response are either dampening down or ramping up. It is often referred to as the “half effect” in Michaelis-Menten reaction terms (22) as it represents the value of *M* (*t*) at which the effect of the term *M/*(*M* + *η*) is at 0.5 (or 50% of its maximal intensity, which is unity). This parameter can only assume values from zero to unity.

Finally, *ϕ* is the phenomenological parameter that aims to capture the “strength” of the illness. This is a very difficult concept that encapsulates many different aspects of the disease and is highly dependent on the patient’s history, pathology, aetiology and other neurological and physiological constructs. Mathematically, it is a parameter similar to the rate of infection in infectious disease modelling: the higher the value, the higher the impact on demyelination and inflammation. In MS disease, microglia, macrophages, T cells and B cells are all known to cause damage to myelin. As such, *ϕ* can be thought of as a proxy for the strength of these cells in their ability to cause damage. Although it is not possible to translate *ϕ* into something immediately measurable, we believe it is reasonable to directly link high values of this parameter to increases in a number of different characteristics of MS’ pathology, including appearance of CELs, heightened EDSS, and increased relapses rates. One of the goals of the present work is to explore how *ϕ* impacts the dynamics of inflammation and demyelination of MS patients.

## 3. STEADY STATES AND LOCAL STABILITY ANALYSIS

To investigate the dynamics and predictive capacity of our system, we first find the model equilibria, which are:

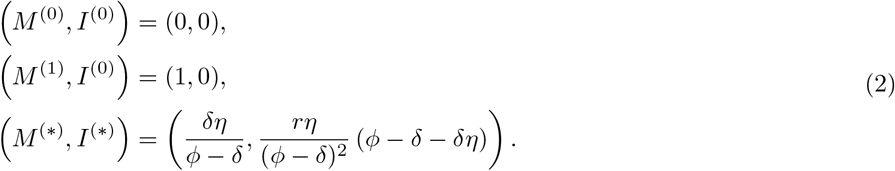

We have a trivial equilibrium (*M* ^(0)^, *I*^(0)^) that could be thought of representing the maximum, unrealistic extent of a patient’s disease: there is no more myelin left and no inflammation is taking place. The other trivial equilibrium (*M* ^(1)^, *I*^(0)^) can be seen as representing an individual who is not affected by demyelination and has no inflammation present. The non-trivial state (*M* ^(*\**)^, *I*^(*\**)^) is the disease equilibrium, where demyelination and inflammation assume positive values for biologically sound parameters. Note that if *ϕ < *δ** or *ϕ < *δ**(1 + *η*) the state (*M* ^(*\**)^, *I*^(*\**)^) is negative and no longer biologically plausible.

Evaluating the Jacobian matrix, we find

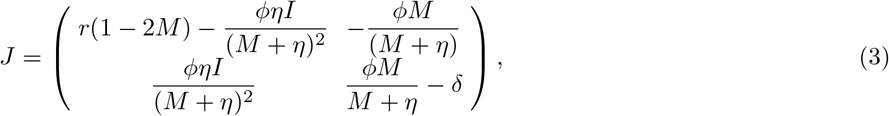

which we use to study the stability of the three steady states. Equilibrium (*M* ^(0)^, *I*^(0)^) is always unstable: the eigenvalues for this steady state are

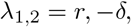

and, for biologically reasonable parameters *r, δ* ∈ *ℝ*_*≥*0_, this is an unstable saddle point. At typical parameter values, the fact that this equilibrium is unstable allows for the system to never go to a state of zero myelin, which would represent an unreasonable state. Because of this, oscillations, when present, are characterised by values of myelination that could be low but never zero.

Evaluating the Jacobian at (*M* ^(1)^, *I*^(0)^) we find that the equilibrium is stable when *ϕ < *δ**(1 + *η*) and unstable otherwise. Note that this is also the condition guaranteeing that *I*^(*\**)^ *>* 0 and, in fact, at *ϕ*^*\**^, i.e. *ϕ* = *δ*(1 + *η*), we have that (*M* ^(*\**)^, *I*^(*\**^)^)^ = (*M* ^(1)^, *I*^(0)^) : as we will see, there is an exchange of stability between the healthy non-diseased steady state and the diseased steady state. This is an interesting mechanism that describes, as we will see shortly, the beginning of a decrease in myelination as the strength *ϕ* of the illness augments.

Lastly, evaluating the Jacobian at (*M* ^(*\**)^, *I*^(*\**)^), we have

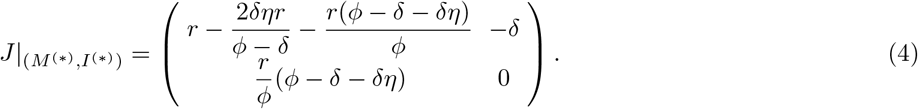

This gives the characteristic polynomial

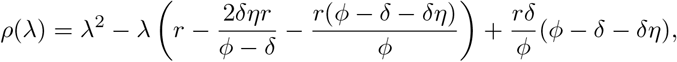

which has roots

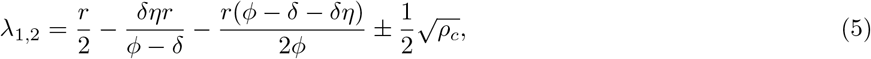

where

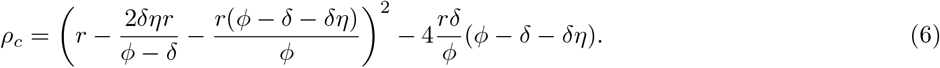

A Hopf bifurcation occurs when Tr(*J*) = 0, det(*J*) *>* 0, and the discriminant is negative, where *J* is the Jacobian evaluated at (*M* ^(*\**)^, *I*^(*\**)^), given by Eq. 4. These conditions give rise to the constraint

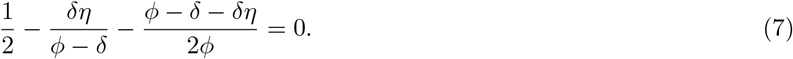

When the Hopf bifurcation occurs, the disease state has transitioned from a stable equilibrium with fixed myelination values to oscillations and periodically changing myelination values. Solving Eq. 7 explicitly yields the Hopf locus:

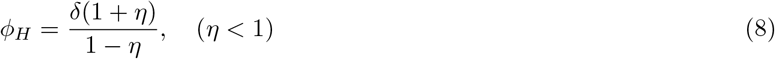

which gives the locus of parameter values for which a Hopf bifurcation occurs and the appearance of limit cycles emerge.

In Fig. 2(B)-(D) we present illustrations of the main dynamics observed from the change in stability of the steady states at (*M* ^(1)^, *I*^(0)^) and (*M* ^(*\**)^, *I*^(*\**)^) . The first example in Fig. 2(B), shows the healthy disease state, when (*M* ^(1)^, *I*^(0)^) is stable. Increasing *ϕ*, we observe an exchange of stability and the diseased steady state becomes stable, see Fig. 2(C). Incrementing *ϕ* further, we pass through the Hopf bifurcation and observe oscillations in myelin and inflammation populations, see Fig. 2(D). In the following section, we provide a bifurcation diagram capturing these dynamics in more detail.

## 4. BIFURCATION ANALYSIS

It is now interesting to understand how the dynamics of the model are affected by changes in its parameters. One of the most compelling is *ϕ*, which represents the severity of MS. We perform a bifurcation analysis using the bifurcation software `AUTO` and `XPPAUT` (15; 17). As shown in Fig. 3(A), the system responds in different ways to an increase in the “magnitude” of the illness. After a value *ϕ*^*\**^ = *δ*(*η* + 1), there is a change in the stability of the healthy equilibrium, which becomes unstable via a branch point (BP). The stable equilibrium shows a progressive decrease in the value of myelination *M*, until it bifurcates into a supercritical Hopf point (HB): limit cycles appear and their amplitude increases up to a maximum, after which they reach a constant magnitude for large values of *ϕ*. All the oscillations that are borne out of a Hopf point in the current model are supercritical: they always emerge as stable limit cycles, with no unstable manifolds and hence a negative first Lyapunov coefficient (27).

**Figure 3.**
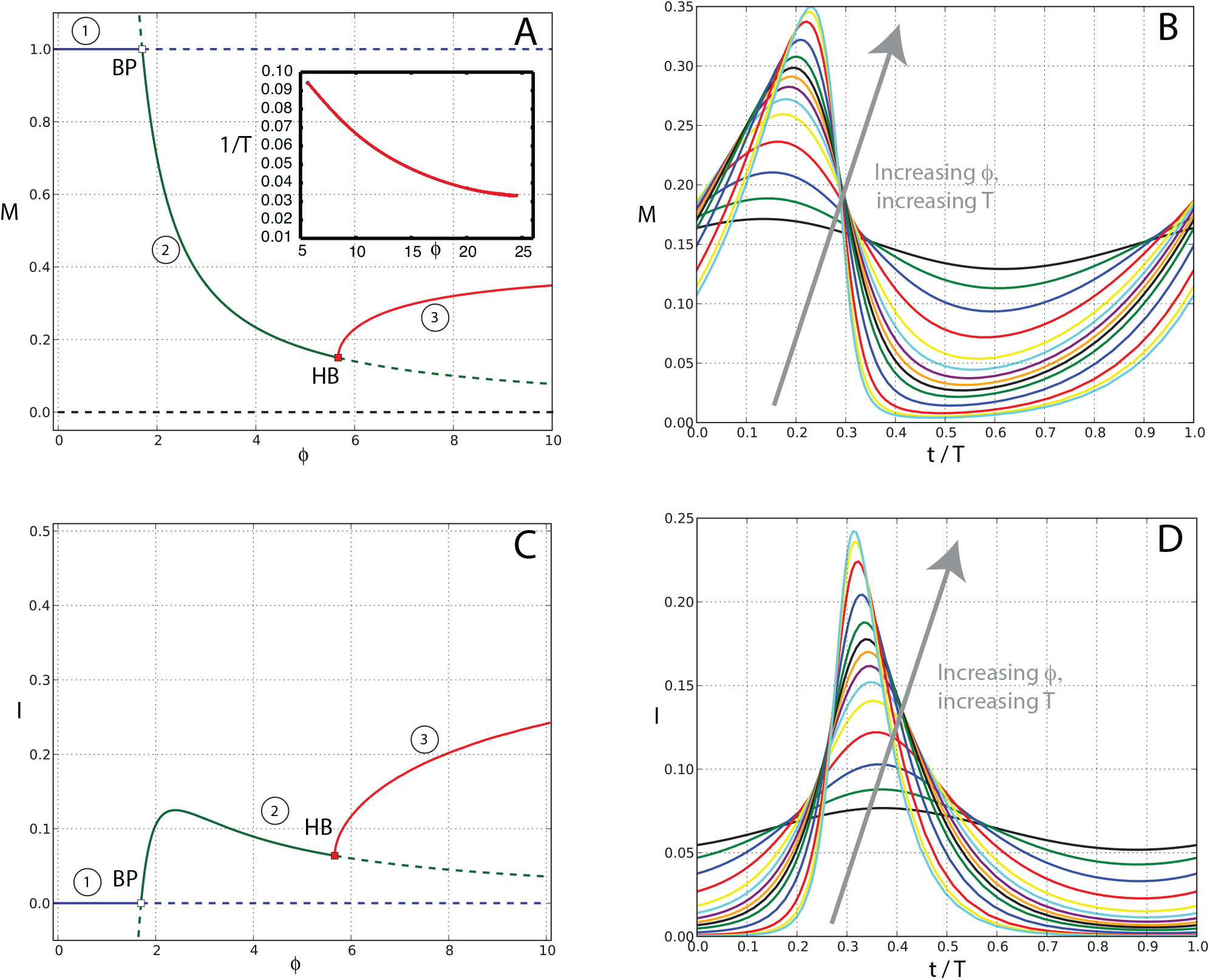
Characteristic bifurcation plots with respect to *ϕ* and periodic orbits originating from the Hopf bifurcations of the model. For (A) and (C), a continuous line indicates a stable equilibrium, a dashed line indicates an unstable equilibrium. HB indicates a Hopf point at *ϕ*_HB_ = 5.67, BP indicates a branch point where stability changes.The value of *ϕ* at which the BP appears is given by *ϕ*^*\**^ = 1.7. Codimension one plots for *M* and *I* are shown in panels (A) and (C) respectively, with the same branches having the same colours. Increasing the strength *ϕ* moves the system from a healthy equilibrium (1) (*M* ^(1)^, *I*^(1)^), to a non-healthy state (2) (*M* ^(*\**)^, *I*^(*\**)^) and finally to a full MS state (3) with limit cycles. Only the maximum amplitude of the cycles is given by the red curve. Oscillations in *M* and *I* for limit cycles are in panels (B) and (D). The period of the oscillations varies from a minimum of about *T* = 10 months to *T* = 20 months. An increasing amplitude corresponds, for the oscillations in this system, to an increasing period or decreasing frequency, as shown in the inset of panel A. Note that all orbits’ periods are normalised to unity (*t/T*), with *T* being the actual period. Other parameters are chosen as *r* = 0.5, *η* = 0.7,*δ* = 1.

These three different phases in the bifurcation diagram seem to mimic one of the possible developments of MS in patients. Starting from an individual with no demyelination, there is then a small steady increase in demyelination, but with no relapses, followed by a final oscillatory state representing relapses of a periodic nature, in line with common symptomatology. The typical shape of such oscillations is presented in Fig. 3(B), with excursions increasing with *ϕ*. Similarly, oscillations’ frequency (respectively, period) also increase (respectively, decrease) with parameter *ϕ*, as shown in the inset of Fig. 3(A). Note that, to emphasise the typical progression of maxima and minima with *ϕ* and other parameters, limit cycles’ period are normalised in all plots throughout this work.

A similar behaviour to myelination is also shown in the inflammation variable *I*, depicted in Fig. 3(C). There, an initially non-inflamed state with *I* = 0 branches (BP) at *ϕ*^*\**^ and the stable equilibrium increases up to a maximal value. The stable steady state branch then starts decreasing until it bifurcates into a Hopf point (HB) and shows oscillations whose maxima increase with *ϕ*, indicating a larger and larger inflammatory response as the strength of MS increases. Note also how the minima of such oscillations occur close to *I* = 0 (see Fig. 3(D)), indicating the presence of remittance events where no (or negligible) inflammation is present. It is interesting to note that, close to HB, the minima of the oscillations are not zero and the orbits have very limited extensions, looking almost constant. So, although periodic, the values of *I* are not dissimilar to those constant values for smaller *ϕ < ϕ*^*\**^. This is, once again, in line with the appearance of an initial phase in the development of relapse-remittance dynamics, where demyelination is still not strong enough to trigger clinically detectable symptoms, and the minima in demyelination values *M* are not low enough to produce a neurologically relevant event.

It is also important to consider how *M* responds to changes in other parameters of the system. For example, in Fig. 4 (A), changes with respect to *η* are depicted, where *η* represents the so-called “half effect” or how “robust” the patient is to inflammation: the higher the *η* the lesser the “impact” of demyelination on the subject. Clearly, when the inflammation affects the patient in a stronger way, *η* decreases and demyelination augments, going from a healthy state to oscillations, which, as previously seen, represent the onset of the disease. The amplitude of the oscillations is much larger in this case (see Fig. 4(B) compared to Fig. 3(B)) and they rise in amplitude very quickly as soon as *η <* 0.5. Qu(ite interes)tingly, as *η →* 0, the trivial equilibrium tends towards the fully diseased equilibrium, i.e. (*M* ^(*\**)^, *I*^(*\**)^) = (*M* ^(0)^, *I*^(0)^) : the origin is unstable (saddle-node) and it turns out that the system allows for solutions that are physiologically irrelevant, with biologically meaningless shapes and periods. This is not surprising, as *η* = 0 represents a value describing the system in a trivial, unrealistic, limit state of zero myelin and zero inflammation.

**Figure 4.**
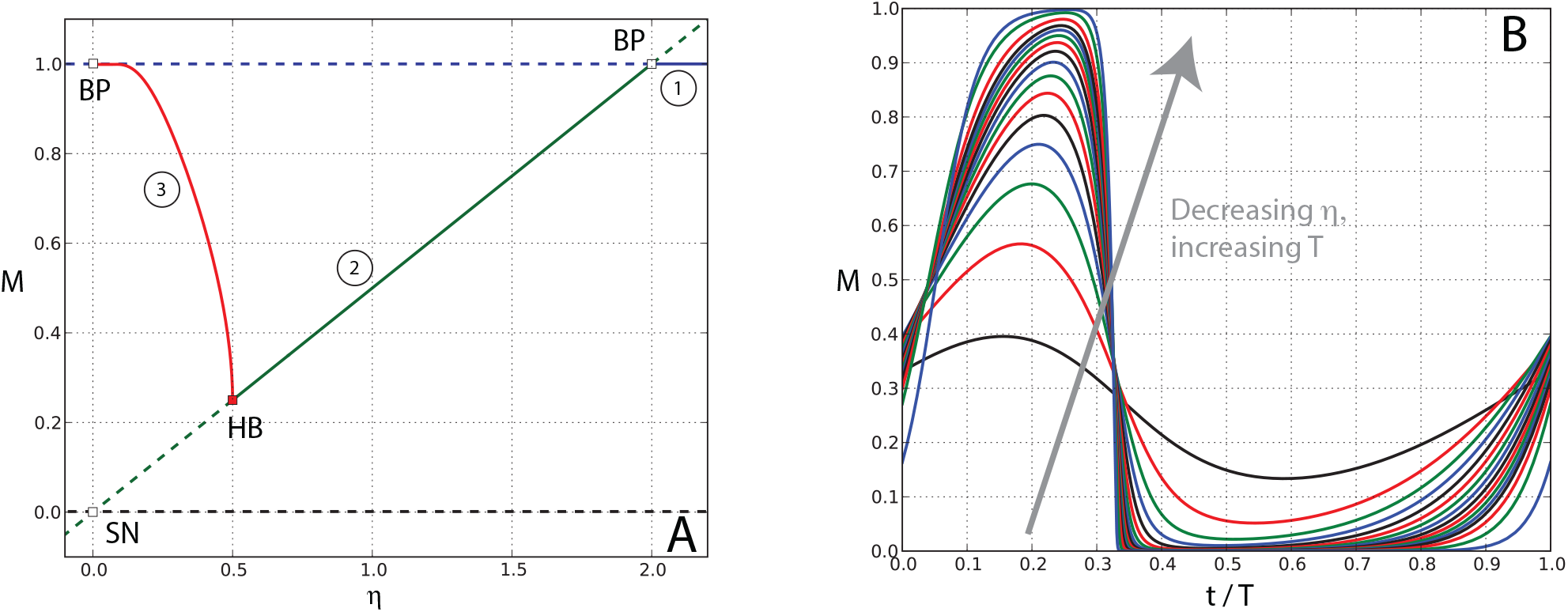
Bifurcation plot with respect to *η* and the periodic orbits originating from the Hopf bifurcation. As per the previous case in Fig. 3, a continuous line indicates a stable equilibrium, a dashed line indicates an unstable equilibrium, HB and BP indicate a Hopf point and a branch point, respectively at *η*^*\**^ = 2.0 and *η*_HB_ = 2.0. Branches in blue and green, indicated with numbers 1 and 2, are loci of stable equilibria, whereas the branch in red, with number 3, pertains to limit point one. Orbits’ periods are normalised to unity. Note that the origin is a saddle-node (SN), because the disease and trivial equilibria coalesce at *η* = 0 but the eigenvalues of this equilibrium are given by *r >* 0 and *− *δ* <* 0. The solution, within the limits of the integration scheme we employ, is a non physiological, almost square-shaped oscillation, with an unrealistically long period. Note that a decreasing *η* leads to limit cycles with a similar sequence (1)-(2)-(3) as in the previous case for *ϕ*. With the parameters we use in this case, since 0 *< η <* 1, only states (2) and (3) are realistic and accessible, whilst state (1) does not have biological significance. We depict results outside this range for illustrative purposes only and to explicitly show the branch change at *η*^*\**^. Parameters are as per the previous bifurcation plot, with *ϕ* = 3.

A similar scenario is present for a codimension one bifurcation plot for *δ* (not shown), which measures the decay rate of the inflammation. In that case, the lower the decay rate, the more prone the system is to produce oscillations, and a similar transition to MS (no illness, decay in myelination, oscillations) is present. It is worth observing that the case *δ* = 0 is also an uncommon, meaningless occurrence and should not be considered as part of the model. The second term of Eqs. 1 in fact reduces to an equation with only the term *ϕMI/*(*M* + *η*) on the right hand side, so that lines of solutions for *M* = 0 and *I* = 0 appear. This case, once again, does not make sense in the context of MS etiology and physiology, and must be discarded as not biologically genuine. It is also relevant to underline that changes in the rate of demyelination *r* do not modify the structure of the bifurcation plots or the shape of oscillations. Nonetheless, a lower *r* generally causes longer transients in the oscillatory dynamics, with the model taking longer time to settle on the limit cycle than the case of a higher *r*..

The importance of oscillations in this model makes the study of the occurrence of Hopf points relevant, as other authors also discuss (50; 51; 16). Eq. (8) gives rise to the plot in Fig. 5(A): this is the portion of parameter space that contains all the HB points in the model. Correspondingly, an example of a section of this surface is presented in Fig. 5(B) as a codimension two plot of the Hopf points. The evidence from these plots is that parameters have different effects on the model dynamics: depending on the rate of decay of inflammation *δ*, the couple (*ϕ, η*) regulates the birth of limit cycles via a Hopf bifurcation in different ways. As *δ* increases, the period of the oscillation borne out of the Hopf point tends to decrease, with small values of (*ϕ, η*) producing oscillations with large periods and vice versa for high values of (*ϕ, η*). Generally speaking, a *T* = 36 months (i.e. 3 years), which can be considered as a maximal value for physiologically meaningful periods, is reached around *ϕ ≈* 0.42, 0.64 and 1.13 for *δ* = 0.1, 0.5 and 1.0 respectively. Any therapeutic intervention that is able to decrease the length of inflammatory events and/or reduce the strength of the MS intensity, will improve patients outcomes. The model can further help clinicians understand how to increase, potentially, the time in between relapses and decrease the amount of relapses for every given year. In fact, in Fig. 6, the loci of points representing limit cycles of the system are depicted according to pairs of given model parameters. For given intensities *ϕ*, the plots show that periods of oscillations tend to increase for increases in *η* and decreases in *δ*, Fig. 6(A),(B) respectively. Given the structure of the orbits for this model, the larger the periods are, the longer the remittance phases. So, any therapeutic approach that can reduce the duration of inflammation or increase *η* can potentially improve patients outcomes by making remittance periods longer. Recall also that *η* measures the ability of the patient to be “resilient” to MS inflammatory events. These data show that there could be dramatic increases in the number months without any relapse, for example, from an average of one relapse every two year to one every four.

**Figure 5.**
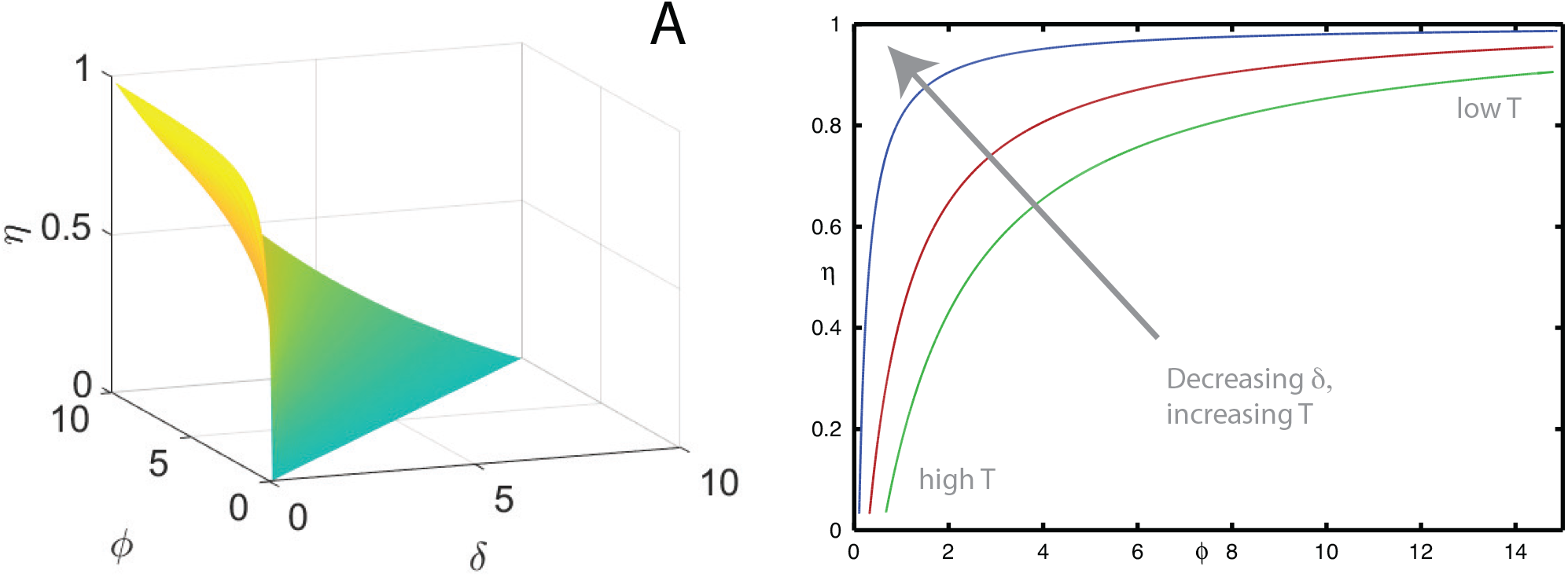
(A) Surface in (*ϕ,δ, η*) space for the set of points satisfying the conditions of the Hopf bifurcation given in Eq. 8. (B) Typical example of a section of the surface at a constant *δ* in the plane (*ϕ, η*), obtained via a codimension two bifurcation plot in such variables. Different branches are obtained for different values of *δ*, i.e. *δ* = 0.1, 0.5 and 1.0 respectively in blue, red and green. Periods of the orbits that are borne out of these Hopf points increase significantly for small *ϕ* and small *η*, whilst, at large *ϕ* and *η*, periods typically range between 9 and 30 months, depending on *δ*. Note that the demyelination rate is kept constant and has value *r* = 0.5.

**Figure 6.**
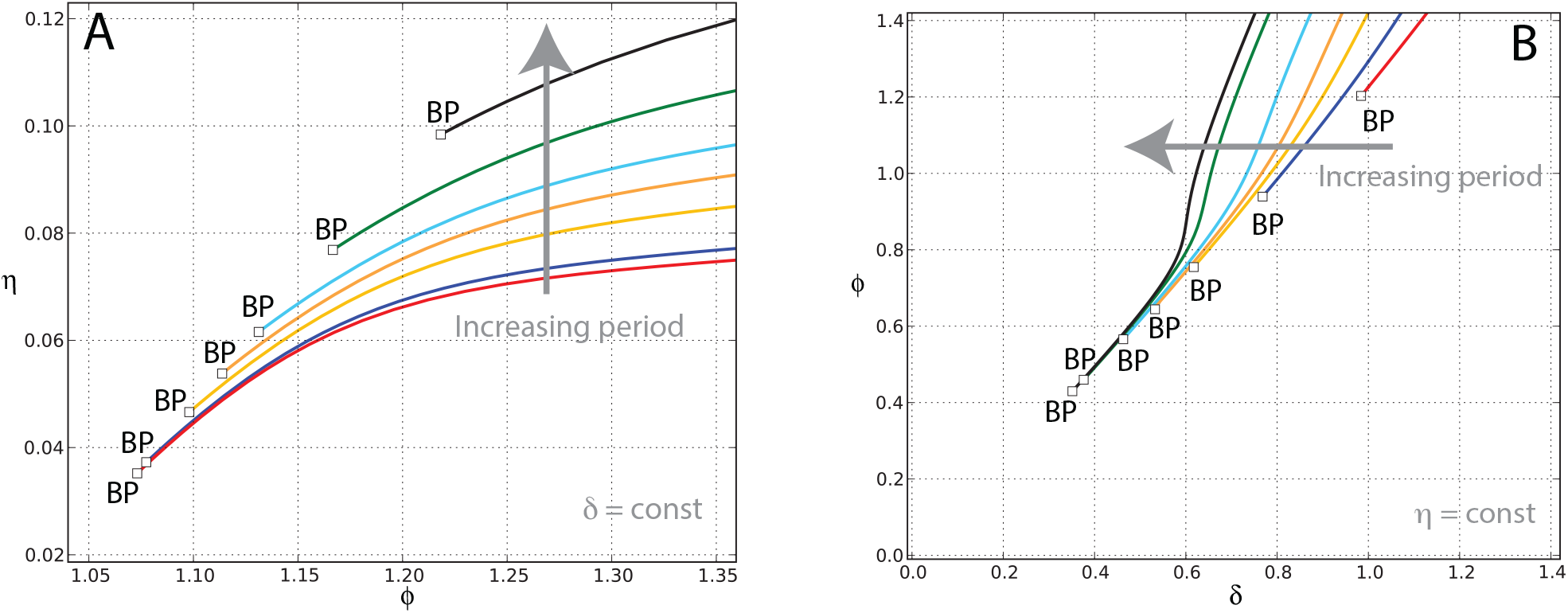
Dependence of oscillation periods on parameters, for different branches of periodic orbits. Each branch has a unique period. The arrow indicates the direction of increase in orbital period, which ranges from about *T* = 20 to *T* = 50 months. BPs (branch points) indicate where the orbits originate (or terminate). Remyelination rate is given by *r* = 0.5 and is the same and constant in both panels. Parameter *δ* is kept also constant in panel (A), whereas panel (B) shows branches for orbits for *η* constant. Note the different values on the axes for the two panels.

## 5. COMPARISON OF MODEL PATTERNS WITH PATIENT DATA

Multiple sclerosis is often characterised by noisy oscillations in patient symptoms and clinical markers. Using the originally published clinical data set from Bagnato *et al*. (2), that was also used by Vélez de Mendizábal *et al*. (46), our model is capable of giving some insight into the types of fluctuations and periodicity that such data show, see Fig. 7. The originally published data is plotted in Fig. 7(A) where CEL, EDSS and relapses were recorded for 9 patients over 45 months (see (2; 46) for further details). Analysing the frequency of peaks in the CEL measurements (Fig. 7(A)), we find that the time between peaks in CELs varies from 2 to 9 months, with an average of 3.7 months. In contrast, peaks in EDSS and relapses are seen much less often, with some patients exhibiting very irregular or spiky dynamics. Some changes to EDSS and relapses occur 18 months apart for some subjects, with a period generally much longer than CEL measurements. This is unsurprising, given that it is established that T1-weighted images show CELs four to ten times more frequently, when compared with clinically defined relapses (46).

**Figure 7.**
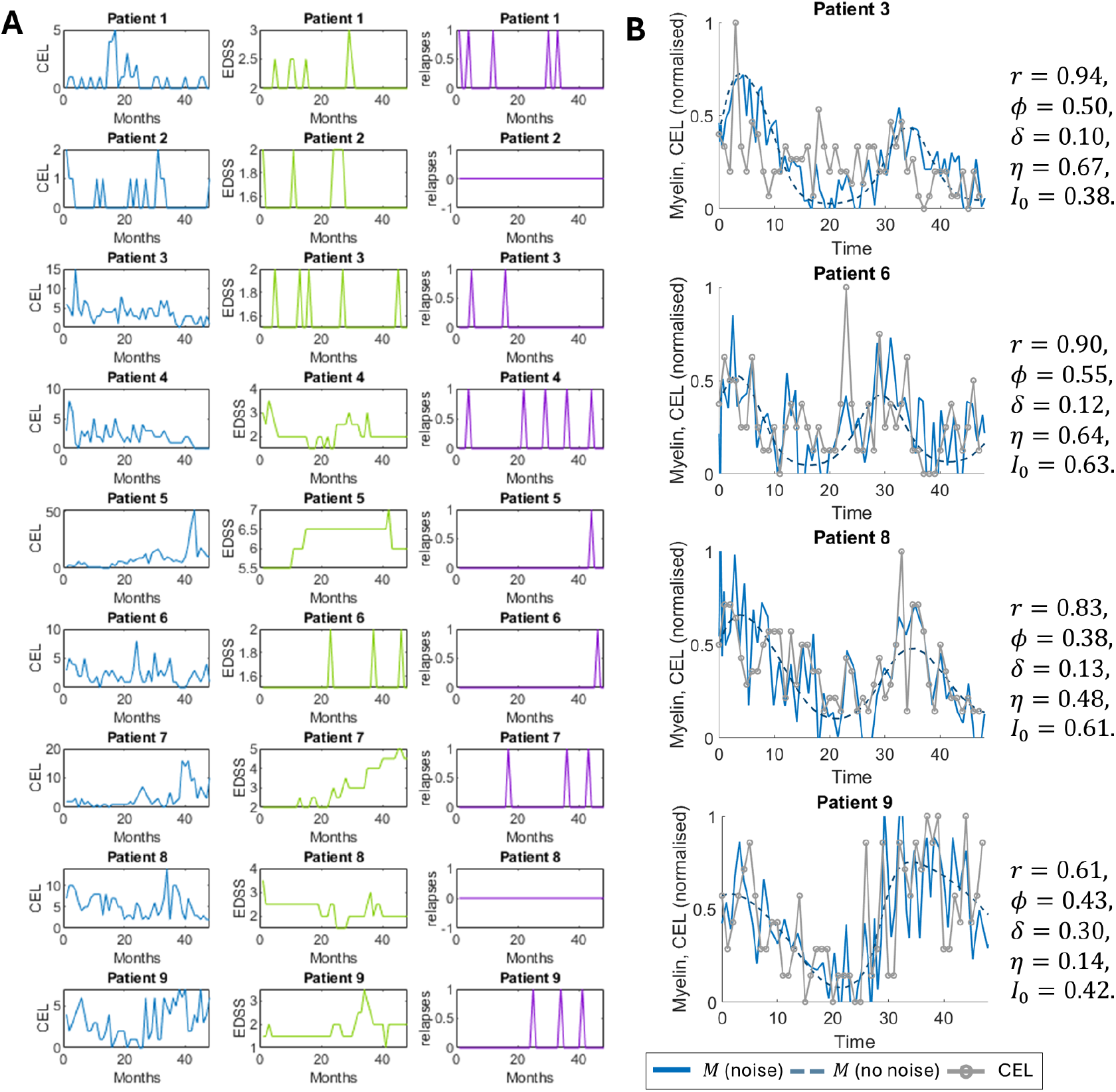
Typical oscillatory dynamics of patient measurements in several biomarkers. (A) Patient measurements over 45 months for contrast enhancing lesions (CELs) (left column), expanded disability status score (EDSS) (middle column) and relapses (right column), originally published in (46; 2). Note the variability across patients and the different structures of the signals. (B) Comparison of the model to CEL measurements for Patient 3, 6, 8 and 9 where the signal has been normalised by the maximum value for each times series, per patient. The model fit (no noise) is given as a dashed line, with an example of the fitted model simulation with Gaussian additive noise, zero mean and 20% standard deviation, given as a solid blue line.

With the hypothesis that inflammation and demyelination detailed in our model can describe CEL measurements, at least from an heuristic and semi-quantitative point of view, we reproduced trajectories in CEL measurements for four patients. First, we select four representative patients who show sufficiently regular signals, such as patient 3, 6, 8 and 9, see Fig. 7(B). After normalising the CEL data by the maximum CEL intensity for each patient, we fit *r, ϕ,δ, η* and *I*_0_ by restricting the search space to the region of limit cycles.. The individual fitted parameter values ranged considerably, for each patient, see Fig. 7(B). We note that the values are similar in their magnitude, but clear differences emerge. For example, Patient 9 has the smallest value of *η*, suggesting their inflammatory response is more saturated than others. Whereas, patient 3 and 6 have the highest rates of remyelination, i.e. *r*. To take into account the noisy nature of the data, we add Gaussian white noise to the model output after estimation of the parameters, centered at zero and with standard deviation equal to 20% of the maximum of the CEL intensity, per patient.

The model is capable of adapting the oscillations to the demyelination data (see dashed curves) and the magnitude of the noise seems appropriate to capture some of the appearance and typical excursions of the fluctuations in the signal. Notably, patient 3 is characterised by a large excursion in real data around 3 months and the signal then almost plateaus for the rest of the series: our system succeeds in describing the trend but only partly and with some shortcomings. Patients 6, 8 and 9 have instead more varied dynamics, with several peaks and troughs throughout, and this results in a better model representation. This is because the dynamical equations proposed here are, in their simplicity, only capable of regular oscillations and, when on a limit cycle, are not able to produce irregular behaviour. Nonetheless, the dynamics of CELs for the patients considered in this example appears to be well represented by this simple model. There are differing conclusions about the link, if any, between CELs and relapses. Given the sparsity of relapses in the data, we chose not to attempt to model this link as the correlation was not strong enough and we did not want to propose a clinical standard without substantive evidence.

## 6. CONCLUSION

Multiple sclerosis is a chronic inflammatory disease characterised by immune-mediated demyelination of neurons in the central nervous system. The number of open questions surrounding MS pathogenesis, aetiology and treatment makes it a challenging and compelling application for mathematical modelling. In this work, we present a possible, minimal way to capture the disease using a system of two ordinary differential equations, with the aim of representing the oscillatory nature of MS and providing a baseline from which other models can be developed and compared.

Through this two-compartment system, we find three main dynamics for MS disease: a healthy disease state, stable MS disease, and oscillating MS disease. Through a bifurcation analysis, we find a key driver of transitioning from a healthy state to a stable disease and then oscillatory disease, is either increases in *ϕ*, i.e. the “strength” of the illness, or decreases in the critical threshold *η* controlling the dampening down or ramping up of the inflammation. This points to two possible mechanisms of targets for therapeutics or biomarkers for monitoring to assess disease progress. Furthermore, we find that the period of oscillations is largely driven by changes in the critical threshold *η* and the decay or clearance of the immune system *δ*. This suggests that if patients experience clear lengthening or shortening in their disease relapse duration, then this is likely due to a shift in their immune systems behaviour and not necessarily the strength of the illness, i.e. *ϕ*.

Clinical guidelines define MS activity as a demonstration of relapse, which can be measured by new or active lesions on MRI or changes to a patient’s EDSS score (35). There are other markers of relapse, such as NFLs, which are seen as a possible future tool to obtain more sensitive measures of underlying disease activity (35). In this work, we present a model that could be used with clinical measurements for CEL to quantify the periodicity of MS myelination and inflammation. We find that the oscillations of the modelling framework can capture noisy oscillations in the real patient data. The model output follows the signal evolution reasonably well and a limited amount of noise is capable of reproducing the typical signals. In its simplicity, the system does show some predictive capability, at least for the cases that display some adequate periodicity. More thorough analysis of the model fitting would need to be performed to make any future disease predictions. This model is a first step in introducing measures of disease activity that can capture inflammatory, neurodegenerative and disability elements, allowing faster identification of underlying disease activity and suboptimal response to therapy. This could further lead to improving risk assessment and treatment decision making (35).

There are some possible amelioration and advances to the model that can be considered. Mathematically, the lack of spatial dependence inhibits us from making predictions on the location of lesions. It could be interesting to formulate a similarly simple approach but consider the non-homogeneous distribution of myelin in different parts of the CNS. This could lead to better predictions of the locations where the lesions appear and disappear, which could be then compared with other seminal works modelling spatial lesion evolution such as Refs. (28; 11). Furthermore, a more in depth characterisation of the immune system, the dynamics of demyelination and the physiology of remyelination could lead to better data fitting and more accurate predictions.

Leading on from this, a limitation of our work is the lack of immunology mechanisms that are modelled. Our hope is that our work could provide simulations to benchmark and calibrate for models that explicitly capture immune dynamics, such as Bisi *et al*. (4). For example, synthetic simulations for inflammation and demyelination could be predicted from our model and compared to the number of effector T cells or activated antigen-presenting cells (APCs) from more detail models of MS immunology. Finally, it could also be interesting to try and better characterise the typical structure of fluctuations in patients’ CEL data. There is a lack of understanding of the causes of such fluctuations and the reasons as to why there is so much variability among subjects.

We hope that the equations presented here can help develop new ideas and methods to tackle MS and eventually improve patient diagnosis and treatment.

